# Restriction of individual branched-chain amino acids has distinct effects on the development and progression of Alzheimer’s disease in 3xTg mice

**DOI:** 10.1101/2025.07.24.663565

**Authors:** Reji Babygirija, Cara L. Green, Michelle M. Sonsalla, Fan Xiao, Mariah F. Calubag, Michaela E. Trautman, Anna Tobon, Ryan Matoska, Chung-Yang Yeh, Isaac Grunow, Diana Vertein, Sophia Schlorf, Bailey A. Knopf, Michael J. Rigby, Luigi Puglielli, Dudley W. Lamming

## Abstract

Dietary protein is a critical regulator of metabolic health and aging in diverse species. Recent discoveries have determined that many benefits of a low protein diet are the result of reduced consumption of the three branched-chain amino acids (BCAAs), leucine, isoleucine, and valine. Intriguingly, each BCAA has distinct physiological and molecular effects, with restriction of isoleucine alone being sufficient to improve metabolic health and extend the lifespan of mice. While restriction of protein or all three BCAAs improves cognition in mouse models of Alzheimer’s disease (AD), the impact of restricting each individual BCAA on the progression and development of AD is unknown. Here, we investigate the effect of restricting each individual BCAA on metabolic health, AD pathology, molecular signaling, and cognition in the 3xTg mouse model. We find that restriction of isoleucine and valine, but not leucine, promotes metabolic health. Restriction of each BCAA had distinct effects on AD pathology and molecular signaling, with transcriptomic analysis of the brain revealing both distinct and shared, and highly sex-specific, molecular impacts of restricting each BCAA. Restricting any of the three BCAAs improved short-term memory in males, with isoleucine restriction having the strongest effect, while restricting valine had the greatest cognitive benefits in females. We identify a set of significantly altered pathways strongly associated with reduced AD pathology and improved cognitive performance in males. Our findings suggest that restricting any of the BCAAs, particularly isoleucine or valine, may form the basis of a novel sex-specific approach to prevent or delay the progression of AD.

## Introduction

Currently, around 6.9 million Americans age 65 and older are living with Alzheimer’s disease (AD) —a number that could double to 13.8 million by 2060 if no significant medical breakthroughs are made to prevent or cure the disease ^1^. While progress has been made in identifying effective interventions, including discovery of monoclonal antibodies that may slow AD progression, these treatments often come with significant side effects, high costs, and limited efficacy, making the search for more precise and effective treatments urgent ^2,3^.

Dietary interventions for disease can be more affordable than costly pharmaceuticals and effective, and calorie restriction (CR) has been shown to slow or prevent the progression of AD in several mouse models ^4–7^. While a CR diet is difficult for most people to adhere to, diets with altered levels of specific macronutrients that do not restrict calories may be easier to follow ^8^. Dietary protein restriction (PR) promotes metabolic health in both humans and mice, and extends lifespan in mice ^9–14^. We recently showed that PR slows progression of AD pathology and preserves cognition in the 3xTg mouse model of AD ^15^.

We previously demonstrated that the metabolic benefits of PR are driven in part by reduced intake of the branched-chain amino acids (BCAAs; leucine, isoleucine, and valine), and showed that restriction of the BCAAs is sufficient to recapitulate the effects of PR on healthspan and lifespan in mice ^9,12^. In subsequent studies, we identified distinct roles for the different BCAAs, showing that restriction of isoleucine is necessary and sufficient for the metabolic benefits of PR, and that isoleucine restriction can extend the lifespan of mice ^16,17^.

In AD mouse models, BCAA restriction has been shown to improve cognitive function, brain AD pathology, and neurotransmitter levels, suggesting a causal link between BCAAs and AD progression ^18^. In the 3xTg mouse model of AD, BCAA supplementation worsens AD neuropathology and reduces survival, whereas BCAA restriction improves cognitive deficits ^19^. However, the role of each individual BCAA in the development and progression of AD remains unknown.

The three BCAAs differentially activate the protein kinase mechanistic target of rapamycin complex 1 (mTORC1), a key regulator of many metabolic processes ^20^. Impaired BCAA metabolism or excessive dietary BCAAs may drive AD pathogenesis by elevating BCAA levels in the brain and activating mTORC1, which is upregulated in the brains of both AD patients and AD mouse models ^19,21–23^. Although BCAA catabolism shares many steps, the intermediate and final products of each BCAA are distinct; they are catabolized either to acetyl-CoA (leucine), propionyl-CoA (valine), or both (isoleucine). Furthermore, there is conflicting evidence regarding individual BCAAs in AD pathogenesis in humans. For instance, analysis of cerebrospinal fluid and plasma amino acid profiles revealed a significant reduction in valine levels in AD patients compared to healthy controls ^24^, while elevated isoleucine levels have been observed in individuals with mild cognitive impairment ^25^.

In this study, we investigated whether the restriction of individual BCAAs could slow or prevent the progression of AD pathology and cognitive loss in the 3xTg mouse model of AD. This model expresses familial human isoforms of APP (APPSwe), Tau (tauP301L), and Presenilin (PS1M146V), and exhibits Aβ and tau pathology in addition to cognitive deficits ^26,27^. We fed 3xTg mice of both sexes either a Control amino acid (AA) defined diet, or diets low in either isoleucine, leucine, or valine starting at 6 months of age which is after the age at which 3xTg mice develop cognitive deficits and aspects of AD pathology. We assessed the effects of restricting each BCAA on metabolic health, AD neuropathology, cognition, and survival. While the effects of each BCAA vary with sex and the specific assay performed, we find generally that restriction of isoleucine and valine, but not restriction of leucine, promotes metabolic health. Isoleucine and valine showed sex-specific effects on both AD pathology and molecular signaling. Restricting any of the three BCAAs improved short-term memory in males, with restriction of isoleucine showing the most pronounced benefits for memory in males; in contrast, restriction of valine showed the greatest benefits for cognition in females. We also found that restricting isoleucine improved the survival of males. Analysis of the brain transcriptome revealed both distinct and shared molecular pathways among all the three individual BCAAs in a highly sex-specific manner. We identified shared and BCAA-specific pathways associated with improved memory and reduced AD pathology. These findings highlight the distinct impact of each BCAA on metabolic health, AD pathology, and cognition, and their sex-specific impact, and suggest that restriction of valine in females or isoleucine in males could serve as a novel approach to prevent or delay the progression of AD.

## Results

### Distinct and sex-specific effects of restricting individual BCAAs on the metabolic health of 3xTg mice

We randomized 6-month-old female and male 3xTg mice to one of four amino acid (AA)- defined diets containing all twenty common AAs; the diet composition of the AA-defined Control (Con) diet reflects that of a natural chow diet in which 21% of calories are derived from protein. The other three diets have a 67% reduction of isoleucine (IleR), leucine (LeuR), or valine (ValR); all diets were isocaloric with identical levels of fat, and the percentage of calories derived from AAs was kept constant by proportionally adjusting the amount of non-essential AAs. These diets, which we have previously utilized ^16^, are detailed in **Supplementary Table 1**. We tracked the mice longitudinally, determining body weight monthly and assessing body composition at the start and end of the study, which spanned nine months. The experimental design is summarized in **Fig. 1A**.

**Figure 1:**
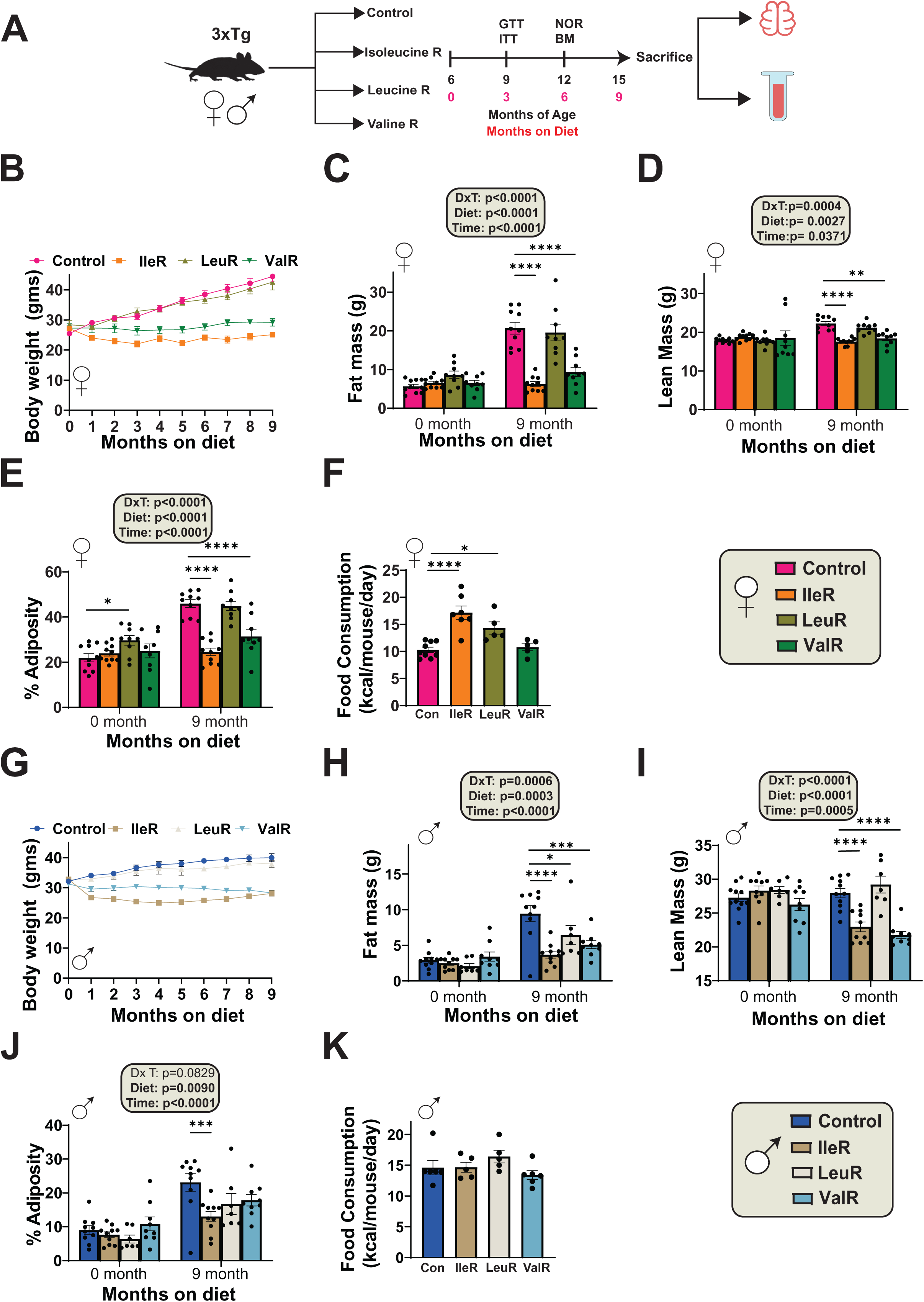
Metabolic health outcomes of 3xTg-AD mice following individual BCAA restrictions. (A) Experimental design: Six-month-old female and male 3xTg-AD mice were placed on amino acid defined Control (Con) diet or in a 67% reduction of isoleucine (IleR), leucine (LeuR), or valine (ValR) and phenotyped over the course of the next 9 months. (B-E) The body weight (B) of female mice was followed over the course of the experiment, fat mass (C) and lean mass (D) was determined at the start and end of the experiment, and the adiposity (E) was calculated. (B-E) n=10 Con, n=10 IleR, n=9 LeuR and n=9 ValR fed 3xTg biologically independent mice. (F) Food consumption of female mice n=9 Con, n=7 IleR, n=5 LeuR and n=5 ValR fed 3xTg biologically independent mice. (G-J) The body weight (G) of male mice was followed over the course of the experiment, fat mass (H) and lean mass (I) was determined at the start and end of the experiment, and the adiposity (J) was calculated. (G-J) n=11 Con, n=10 IleR, n=7 LeuR and n=9 ValR 3xTg biologically independent mice. (K) Food consumption of male n=7 Con, n=5 IleR, n=5 LeuR and n=6ValR 3xTg biologically independent mice. (C-E, H-J) Statistics for the overall effect of diet and time represent the p value from a two-way analysis of variance (ANOVA); *p<0.05, statistics for the overall effects of diet and time represent the p value from a 2-way ANOVA conducted separately for each time point; *p<0.05, **p<0.01, ***p<0.001, **** p<0.0001 from a Dunnett’s post-test examining the effect of parameters identified as significant in the 2-way ANOVA.. (F, K) *p<0.05, **** p<0.0001 one-way ANOVA, followed by a Dunnett’s. Data represented as mean ± SEM.

Both IleR-fed and ValR-fed female 3xTg mice maintained their body weight over the course of the study, while Control-fed and LeuR-fed females continued to gain weight (**Fig. 1B**). IleR- and ValR-fed mice exhibited reduced fat mass and lean mass accretion during the 9-month study; by the end of the experiment, we observed an overall effect of diet on both lean and fat mass (**Fig. 1C-D**). Thus, by the completion of the study, IleR- and ValR-fed females had reduced adiposity compared to Control-fed females (**Fig. 1E**). These changes in body weight and body composition were not the result of reduced caloric intake; rather, as we have previously observed in other mouse strains, IleR-fed females consumed more food than Control-fed females (**Fig. 1F**)^17^. We observed increased food consumption in LeuR-fed mice as well, while ValR-fed females did not eat less.

The result of restricting the individual BCAAs on the weight and body composition of male 3xTg mice were similar, except that LeuR-fed males had reduced accretion of fat mass and an overall reduction of adiposity (**Figs. 1G-J**). Surprisingly, and unlike in females, we did not observe a significant effect of any of the BCAAs on food consumption (**Fig. 1K**).

Since the mice on IleR and ValR restricted diets gained less weight than Control-fed mice despite similar or increased food consumption, we examined the energy balance of all groups using metabolic chambers. We assessed substrate utilization by examining the respiratory exchange ratio (RER), which is calculated using the ratio of O_2_ consumed and CO_2_ produced; the RER approaches 1.0 when carbohydrates are primarily utilized for energy production and approaches 0.7 when lipids are the predominant energy source ^28,29^.

We expected based on our previous findings that mice fed the IleR and ValR diets would have increased energy expenditure ^16^. Indeed, we found that a ValR diet significantly increased energy expenditure in 3xTg females, while IleR-fed mice showed a non-significant increase (p=0.1614, dark cycle) (**Fig. 2A**). No diet-induced differences in activity level or RER were observed (**Figs. 2B-C**). The results in males were similar, with IleR but not ValR significantly increasing energy expenditure; there was also decreased activity in IleR-fed and ValR-fed males, without significant effects on RER (**Figs. 2D-F**).

**Figure 2:**
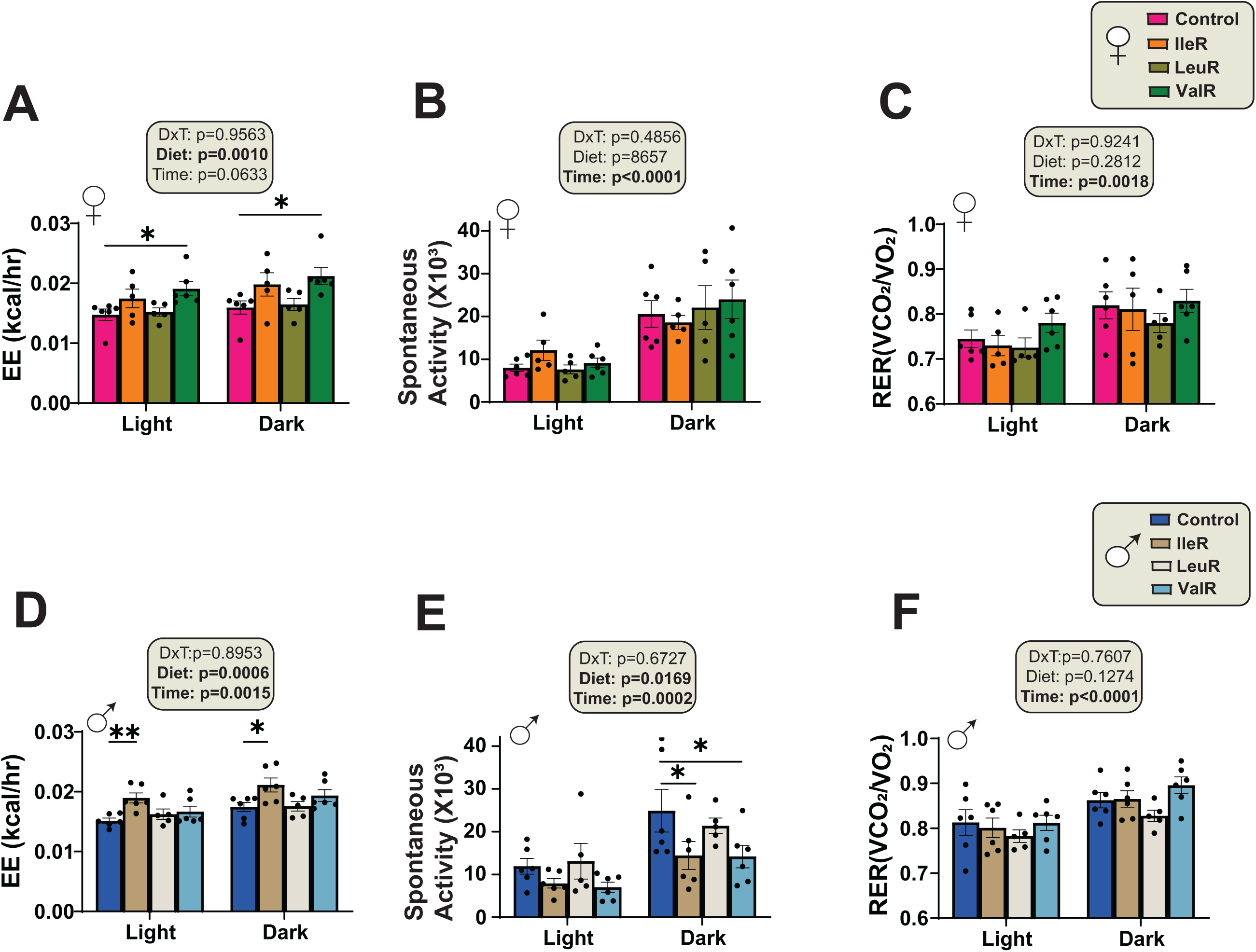
Distinct effects of individual BCAA restriction on energy balance in 3xTg mice. (A-F) Metabolic chambers were used to determine energy expenditure, spontaneous activity and fuel source utilization, over 24 hours in female (A-C) and male (D-F) six-month-old 3xTg fed control, IleR, LeuR or ValR diets for 3 months. (A, D) Energy expenditure normalized to body weight in females (A) and males (D). (B, E) Spontaneous activity of females (B) and males (E). (C, F) Respiratory exchange ratio (RER) in females (C) and males (F). (A-C) n=6 Con, n=5 IleR, n=5 LeuR and n=6 ValR fed 3xTg biologically independent mice. (D-F) n=6 Con, n=6 IleR, n=5 LeuR and n=6 ValR fed 3xTg biologically independent mice. (A-F) Statistics for the overall effect of diet and time represent the p value from a two-way ANOVA; *p<0.05, **p<0.01 Dunnett’s post-test examining the effect of parameters identified as significant in the two-way ANOVA. Data represented as mean ± SEM.

We have previously shown that 3xTg mice have impaired glucose tolerance that is improved by PR in both females and males ^15^. As such, we assessed the effect of restricting each BCAA on glycemic control by performing glucose (GTT) and insulin (ITT) tolerance tests at approximately 9 months of age, after the mice had been on their respective diets for about three months. In female 3xTg mice, IleR feeding improved glucose tolerance (**Fig. 3A**); interestingly, a ValR diet impaired insulin sensitivity (**Fig. 3B**). In males, a IleR diet also improved glucose tolerance, but there was no impact of diet on insulin sensitivity (**Figs. 3C-D**).

**Figure 3:**
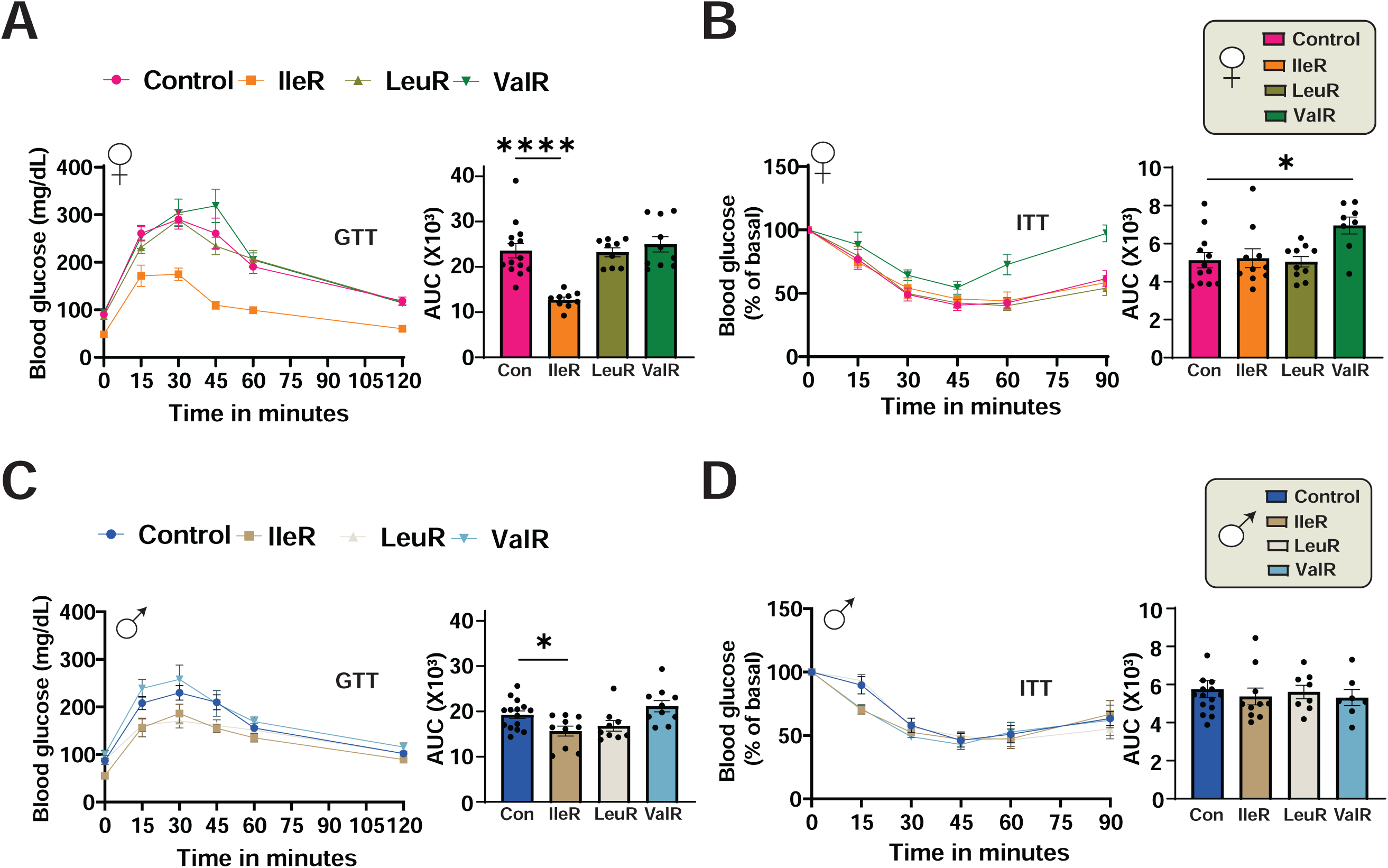
Isoleucine restriction improves glucose tolerance in 3xTg-AD mice of both sexes. (A-B) Glucose (A) and insulin (B) tolerance tests were performed in female mice after three months on Con, IleR, LeuR and ValR fed diets. (A) GTT: n=14 Con, n=9 IleR, n=9 LeuR and n=10 ValR 3xTg biologically independent mice per group. (B) ITT: n=12 Con, n=9 IleR, n=9LeuR and n=9 ValR 3xTg biologically independent mice per group. (C-D) Glucose (C) and insulin (D) tolerance tests were performed in male mice after three months on Con, IleR, LeuR and ValR fed diets. (A) GTT: n=14 Con, n=10 IleR, n=9 LeuR and n=10 ValR fed 3xTg biologically independent mice (B) ITT: n=14 Con, n=10 IleR, n=9 LeuR and n=8 ValR fed biologically independent mice. (A-D) Dunnett’s post-test examining the effect of parameters identified as significant in the one-way ANOVA. *p < 0.05, ****p < 0.0001 Data represented as mean ± SEM. AUC, Area Under the Curve.

### Restriction of specific BCAAs improves neuropathology in 3xTg mice

We next assessed the effect of individual restriction of each BCAA on the progression of AD neuropathology by evaluating several pathological hallmarks of AD, including amyloid beta (Aβ) plaque deposition, phosphorylation of tau, and gliosis. Based on our previous characterization of this model ^15^, we examined AD neuropathology in 15-month-old female and male 3xTg mice, after 9 months of consuming either the Control or individual BCAA restricted diets.

We observed substantial Aβ plaque accumulation in the hippocampus of Control-fed females; IleR-fed and LeuR-fed mice had substantially reduced plaque deposition, while ValR- fed mice actually had double the number of plaques as Control-fed females (**Fig. 4A**). Fluorescent immunostaining and quantitative analysis of hippocampal phosphorylated tau (p-Tau Thr231) revealed that IleR and LeuR-fed females had significantly reduced Tau phosphorylation (**Fig. 4B**). At the level of the whole brain, we observed a significant reduction in p-Tau Thr231 only in ValR- fed females (**Fig. S1A**). Finally, neuroinflammation is a key pathological feature of AD, and we assessed activation of astrocytes and microglia. We conducted immunostaining of brain sections with anti-glial fibrillary acidic protein (GFAP), an astrocyte marker, and with anti-ionized calcium binding adaptor molecule 1 (IBA-1), a microglia marker. While there was no effect of diet on astrocytic activation, restriction of any BCAA reduced microglial activation, reaching significance in the case of IleR- and ValR-fed females (**Fig. 4C**).

**Figure 4:**
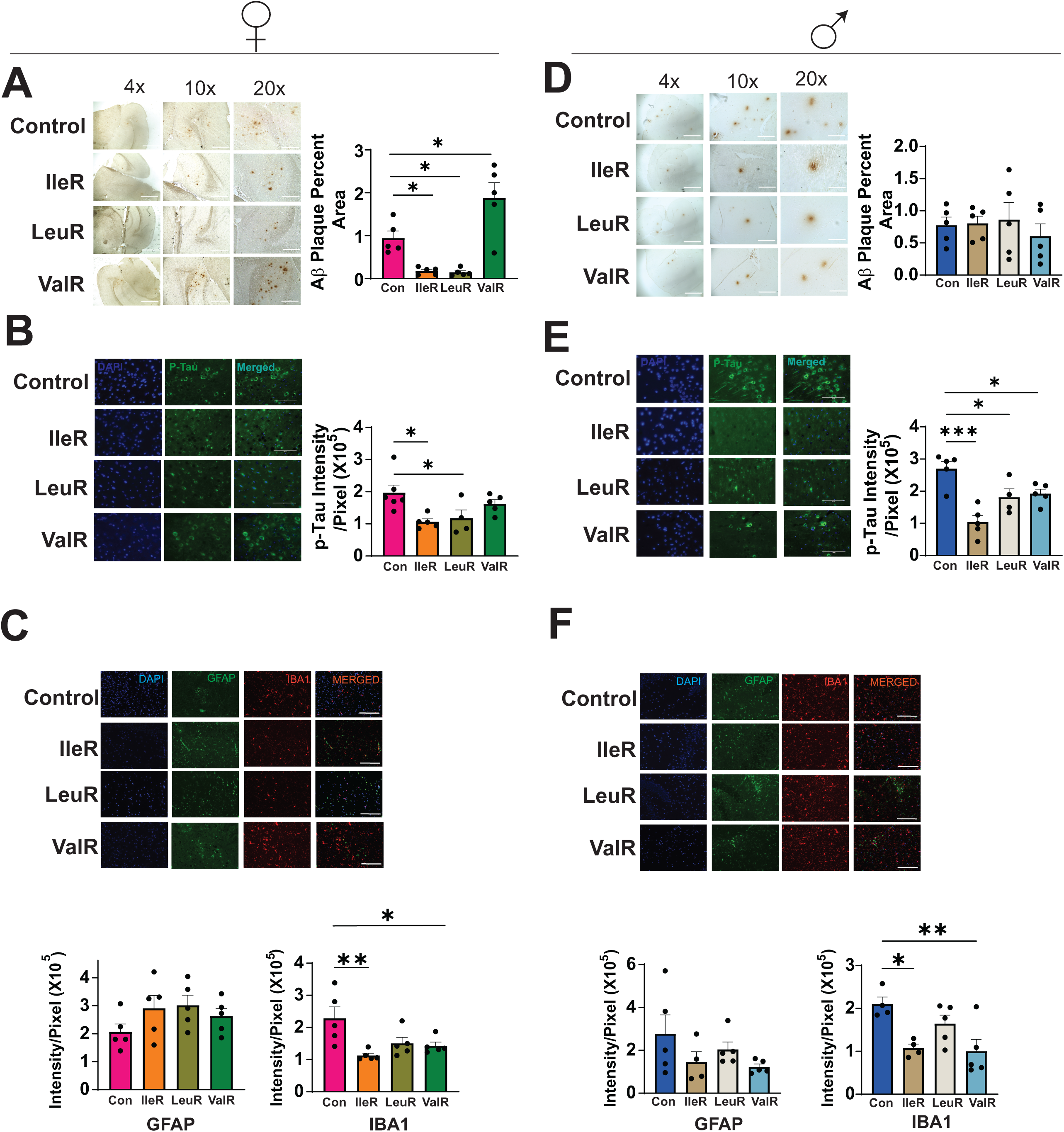
Restriction of isoleucine and leucine improves neuropathological outcomes in a sex dependent manner in 3xTg-AD mice. (A-F) Analysis of AD neuropathology in female and male 3xTg mice fed on Con, IleR, LeuR or ValR diets from 6-15 months of age. (A, D) Representative plaque images of DAB staining with 6E10 antibody in the hippocampus of female (A) and male (D) 3xTg mice. 4x, 10x, 20x and 40x magnification shown; scale bar in the 4x image is 1000 µM, 10x image is 400µM, 20x is 200 µM and 40x is 100 µM. Quantification of plaque area in females shown under representative blots from hippocampus. (A, D) For hippocampus n=4-5 3xTg biologically independent mice per group were used in both sexes. (B, E) Representative immunofluorescence images in the hippocampus of 3xTg females (B) and males (E) stained with p-Tau Thr231 antibody (AT180), 40x magnification shown; scale bar 100 µM. Quantitative analysis of fluorescence intensity. n=4-5 3xTg biologically independent mice per group. (C, F) Immunostaining and quantification of 5 μm paraffin-embedded brain slices for astrocytes (GFAP) and microglia (Iba1) in female 3xTg mice (C) and male mice (F). 20x magnification shown; Scale bar is 200 µM. n=4-5 biologically independent mice/group. (A-F) *p<0.05, ** p<0.01, ***p<0.001 Dunnett’s post-test examining the effect of parameters identified as significant in the one-way ANOVA. Data represented as mean ± SEM.

In 3xTg males, Aβ plaque deposition was less prominent than in females, and there was no effect of restricting any of the BCAAs (**Fig. 4D**). However, restriction of any BCAA in males had significantly reduced hippocampal p-Tau Thr231 as shown by fluorescent immunostaining (**Fig. 4E**). At the whole brain level, there was no effect of IleR or LeuR diets on p-Tau, but ValR- fed males had a significant increase in p-Tau (**Fig. S1B**). While there was no effect of diet on astrocytic activation, restriction of either Ile or Val, but not Leu, significantly reduced microglial activation in the hippocampus, as shown by reduced IBA-1 expression (**Fig. 4F**).

### Transcriptomic profiling of the brain reveals sex-specific and overlapping molecular responses to individual BCAA restriction in 3xTg mice

To obtain insight into the molecular pathways altered by individual BCAA restriction, we next performed transcriptomic analysis of the whole brain of both male and female 3xTg mice fed the Control, IleR, LeuR or ValR diets. We identified the differentially expressed genes (DEGs) expressed as a result of restricting each of the BCAAs. Interestingly, we identified a shared set of 1,623 DEGs genes that were differentially expressed across all the three BCAA-restricted groups in 3xTg males (**Fig. 5A and Table S2**). Restriction of each individual BCAA also resulted in DEGs that were unique to that group, with ValR-fed males showing the largest number (1,688) of DEGs vs. Control-fed males that were not shared with either LeuR or IleR-fed males. We failed to observe any DEGs shared across the three restricted diets in female 3xTg mice (**Fig. S2 and Table S3**).

**Figure 5:**
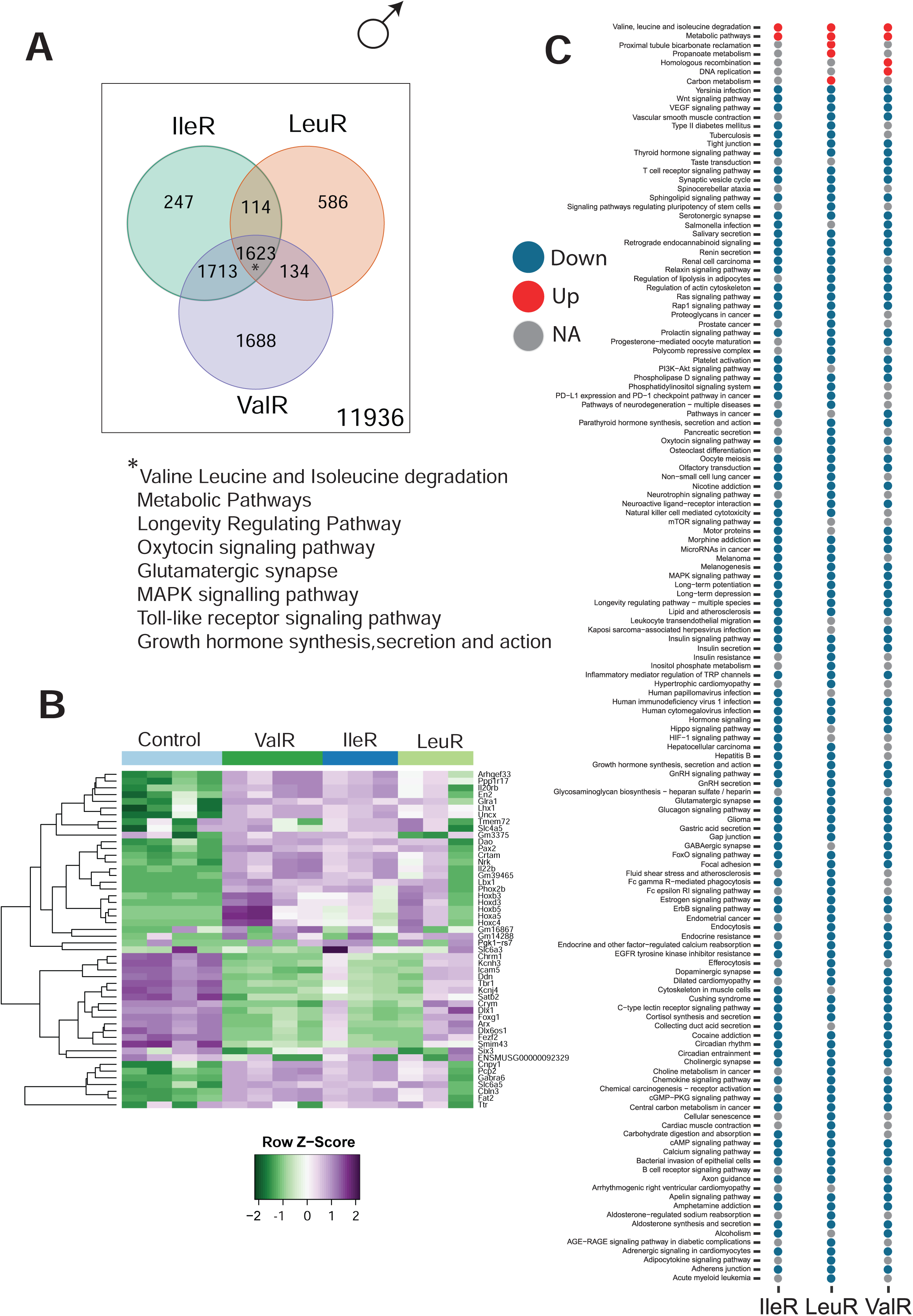
Transcriptional profiling of brain identifies overlapping molecular responses to individual BCAA restriction in male 3xTg mice. (A) Venn diagram of the differentially expressed genes in the three BCAA restricted groups. * The pathways listed below the Venn diagram represent a subset of significantly enriched KEGG pathways (p<0.05) from differentially expressed genes altered by all three BCAA restricted diets. (B) Heatmap of the top 50 differentially expressed (DEG) genes (C) Enriched transcriptomic pathways across all male groups (red = upregulated, blue = downregulated, grey = not significant). n=4-5 mice/group.

Analyzing the expression of the 50 most variable genes revealed shared and distinct gene expression patterns across diet groups. Among the differentially expressed genes, several Hox homeobox genes (*Hoxb3, Hoxa5, Hoxc4, Lhx1*), which have been shown to play critical roles in neurodevelopment and synaptic plasticity and have been implicated in AD pathology ^30,31^, showed altered expression across all the restricted diets in male 3xTg mice (**Fig. 5B**). Consistent with the distinct gene expression patterns observed across all the individual BCAA diet groups; we also found that *Foxg1* (Forkhead box G1), which has been implicated in neuronal development ^32^, was downregulated in ValR and IleR-fed males; however expression in LeuR-fed males was similar to that of Control-fed males. Furthermore, we found that *Satb2,* a gene known for its role in neurodevelopment and cognition, was downregulated in both ValR and IleR-fed males^33^. We also found that *Crym*, which encodes the neuroprotective protein µ-crystallin, was downregulated in ValR and IleR-fed males ^34,35^.

We next performed pathway enrichment analysis to identify biological pathways altered in response to BCAA restriction. In male 3xTg mice, we observed substantial overlap in the pathways across all three restricted BCAA groups, with most pathways being downregulated in two or more groups (**Fig. 5C** and Table S4). Although most of the pathways affected by BCAA restriction were downregulated, a few were upregulated, with “Valine, Leucine, and Isoleucine Degradation” and “Metabolic Pathways” consistently upregulated in all of the restricted groups (**Fig. 5C**). These upregulated pathways could reflect the compensatory metabolic adaptation in the brain occurring due to the limited intake of BCAAs. “Carbon Metabolism & Propanoate Metabolism” was upregulated exclusively in LeuR mice, perhaps suggesting the increased need for alternate carbon sources to meet the metabolic demands caused due to leucine restriction. “DNA Replication” was selectively upregulated in ValR-fed mice, which could suggest increased cellular activity or stress response, possibly reflecting reactive glial cell activation or DNA repair mechanisms ^36^. As ValR males also exhibited worsened AD pathology, specifically increased p-tau in whole brain, this change might represent a pathological cellular response.

Most overlapping pathways were downregulated. Neuroinflammatory pathways were downregulated across all three BCAA-restricted groups, including “MAPK signaling pathway” and “Toll-like receptor signaling pathway,” potentially contributing to the observed cognitive improvements and reduced microglial activation. In addition, the “Longevity-regulating pathway” and “Growth hormone synthesis and secretion” pathways were downregulated in all three restricted groups, suggesting a potential reduction growth and aging-related signaling. It was particularly interesting to see down regulation of “mTOR signaling pathway” only in the IleR group, as leucine is a very potent mTOR agonist. We also observed LeuR-specific downregulation of pathways, including “pathways in neurodegeneration” and “cellular senescence pathways”. In contrast, “Pathways in cancer” was downregulated only in IleR and ValR mice.

Together, these findings indicate that while individual BCAA restriction induce distinct molecular effects, they also engage overlapping molecular mechanisms in males. These shared molecular mechanisms regulate pathways involved in neuroinflammation as well as metabolism and likely contribute to the ability of individual BCAA restriction to delay the progression of AD pathology in males.

### Isoleucine restriction downregulates autophagy and reduces mTORC1 activity in 3xTg mice

We were surprised that autophagy was not implicated as an altered pathway in our transcriptional analysis, and that mTOR signaling was identified as altered only in IleR-fed mice. Both autophagy and mTOR are heavily implicated in the pathogenesis of AD; impaired autophagy machinery leads to ineffective clearance of plaques and tangles, contributing to the progression of AD symptoms in both patients and animal models ^37–41^, while hyperactivation of mTOR in AD disrupts the proteostasis network and acts to inhibit autophagy, exacerbating the accumulation of plaques and tangles ^42^. Inhibiting mTOR, either pharmacologically with the drug rapamycin or via feeding of a PR diet, promotes autophagy and reduces AD pathology ^15,43–46^. We therefore further investigated the effect of individual BCAA restriction on autophagy and mTOR signaling at the protein level.

We performed immunoblotting on brain lysates to assess several autophagy markers, including autophagy proteins ATG5, ATG7, and ATG16, as well as autophagosome formation proteins Beclin and light chain 3A/B (LC3A/B), and the autophagy receptor p62 (sequestosome 1, SQSTM1). In female 3xTg mice, IleR-fed mice exhibited significantly reduced expression of ATG5, ATG7, LC3A/B, and p62 compared to Control-fed mice. LeuR and ValR-fed females showed a significant decrease in LC3A/B expression, a key marker of autophagosome formation, but no significant changes in any other markers (**Figs. 6A-G**). In 3xTg males, we observed a significant downregulation of Beclin in all three BCAA-restricted groups compared to control-fed mice. Although certain other autophagy markers exhibited a decreasing trend in IleR and ValR- fed mice, none of these changes reached statistical significance (**Figs. 6H-N**).

**Figure 6:**
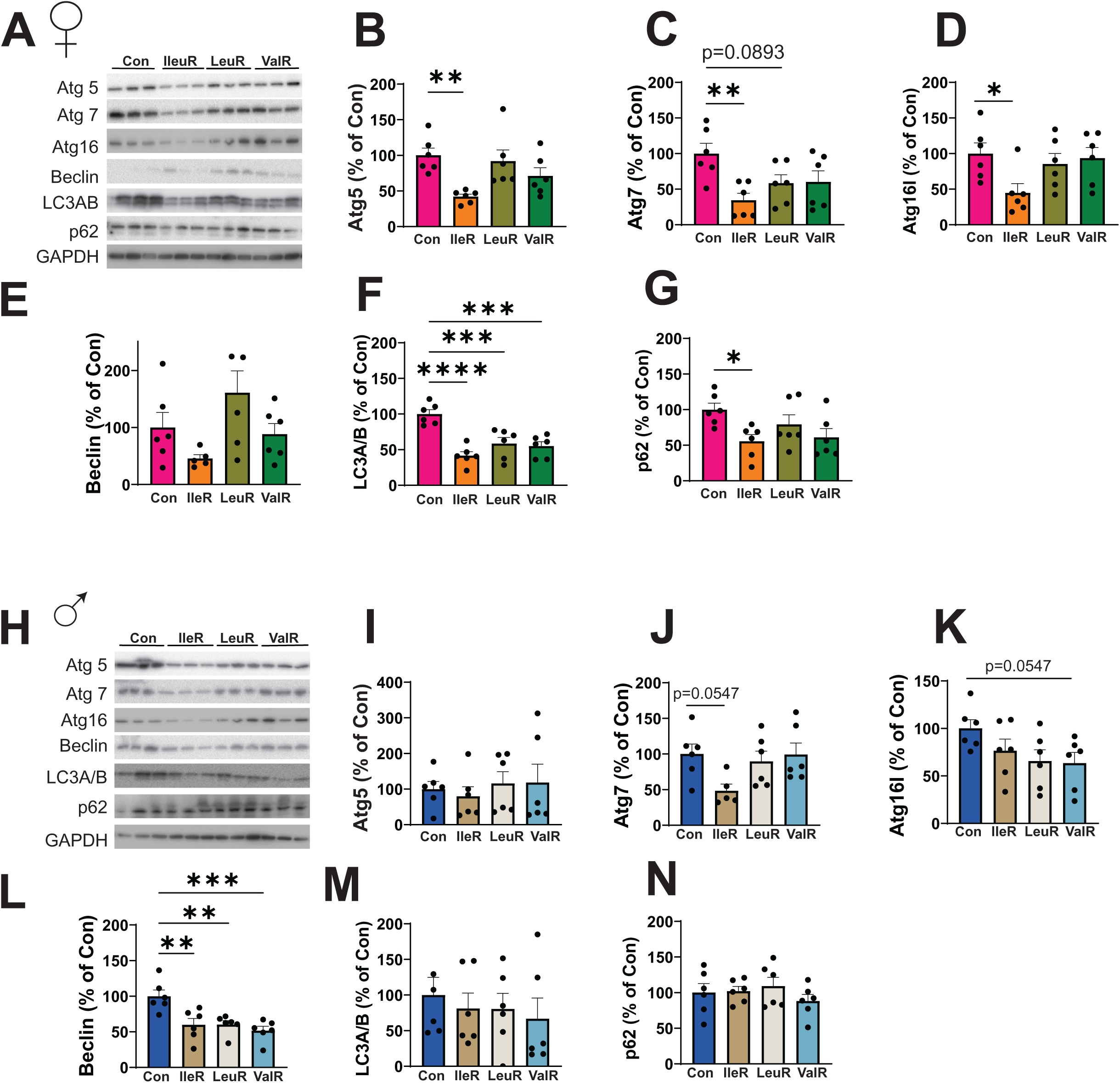
Distinct effects of individual BCAAs on autophagy in 3xTg-AD mice. (A-N) Immunoblotting on brain lysates of 15-month-old 3xTg mice fed on Con, IleR, LeuR or ValR mice to assess several autophagy markers, including autophagy proteins ATG5, ATG7, and ATG16, as well as autophagosome formation proteins Beclin and light chain 3A/B (LC3A/B), and the autophagy receptor p62 (sequestosome 1, SQSTM1). (A) Representatives immunoblot of all the autophagy related proteins in females. (B-G) Quantification of ATG5 expression (B), ATG7 (C), ATG16 (D), Beclin (E) LC3A/B (F) and p62 (G) relative to expression of GAPDH in females (B-G) n=6 3xTg biologically independent mice per group. (H) Representative immunoblot of all the autophagy related protein in males. (I-N) Quantification of ATG5 expression (I), ATG7 (J), ATG16 (K), Beclin (L) LC3A/B (M) and p62 (N) relative to expression of GAPDH in males (I-N) n=6 3xTg biologically independent mice per group. (B-G, I-N) *p<0.05, **p<0.01, ***p<0.001, ****p<0.0001 Dunnett’s post-test examining the effect of parameters identified as significant in the one-way ANOVA. Data represented as mean ± SEM.

We next evaluated mTORC1 activity by performing immunoblotting for the phosphorylation of its substrates, p-S240/S244 S6 and T37/S46 4E-BP1, in whole brain lysates of both female and male mice (**Figs. S2A-E**). In female 3xTg mice, none of the individual BCAA restricted diets led to significant changes in phosphorylation of S240/S244 S6 compared to the Control-fed mice. However, phosphorylation of T37/S46 4E-BP1 was significantly reduced in IleR- fed and LeuR-fed 3xTg females compared to Control-fed mice (**Figs. S2A-C**). In 3xTg males, restriction of any of the three BCAAs resulted in reduced phosphorylation of p-S240/S244 S6 as compared to Control-fed mice (**Figs. S2D-E**). However, while ValR-fed 3xTg males had reduced phosphorylation of T37/S46 4E-BP1 relative to Control-fed mice, IleR-fed 3xTg males unexpectedly had increased T37/S46 4E-BP1 phosphorylation compared to Control-fed mice (**Fig. S2F**).

### Sex-specific benefits of individual BCAA restriction on hippocampal-dependent spatial learning-associated memory deficits

To evaluate the effects of individual BCAA restriction on cognition, we conducted behavioral assays on 12-month-old 3xTg mice fed either a Control, IleR, LeuR, or ValR diet. We examined the performance of the animals in a Barnes Maze and tested Novel Object Recognition (NOR).

In the Barnes maze, mice were required to locate an escape box placed at the target hole using spatial cues learned during a four-day acquisition phase. Short-term memory (STM) and long-term memory (LTM) were tested on days 5 and 12, respectively. Among the female 3xTg mice, ValR-fed females located the escape box the fastest during the training phase and showed the best performance during both the STM and LTM tests, taking significantly less time to identify the escape hole during the STM test (**Fig. 7A**). No significant differences were observed in error hole visits across all groups during training sessions or the STM and LTM tests, but consistent with their improved latency, ValR-fed females tended to make fewer errors than mice fed the Con diet (**Fig. 7B**).

**Figure 7:**
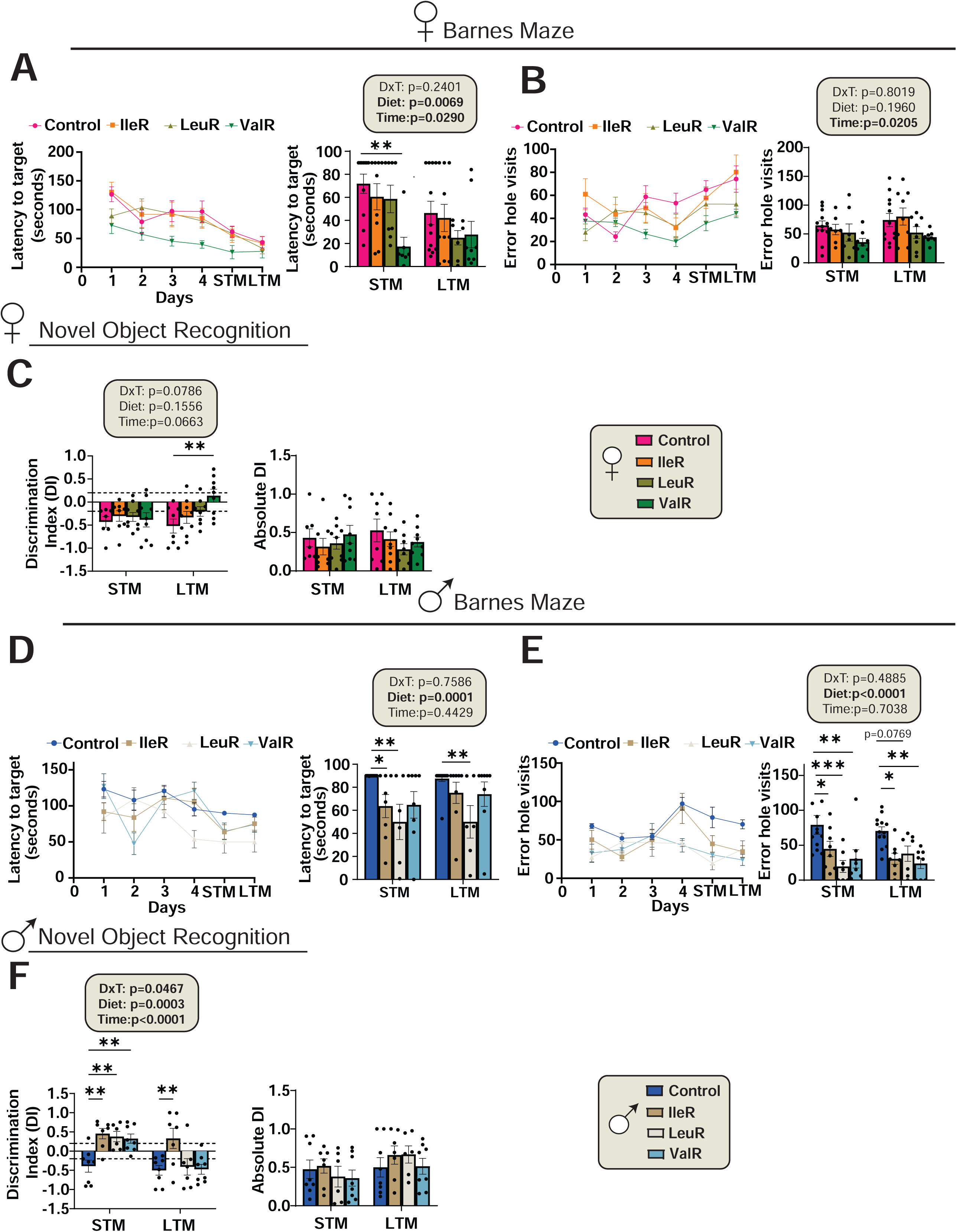
Valine restriction improves cognition in females, while Isoleucine and Leucine restriction improves memory in males. (A-F) The behavior of female and male 3xTg-AD mice was examined at 12 months of age after mice were fed the indicated diets for 6 months. (A, D) Latency of target in Barnes Maze acquisition period over the five days of training and in short term memory (STM) and long-term memory (LTM) tests in female (A) and male mice (D). (B, E) The number of error hole visits during Barnes maze training phase in STM and LTM tests by female (B) and male (E) mice. (C, F) The preference for a novel object over a familiar object was assayed in female (C) and male (F) mice via STM and LTM tests. The dashed lines at +0.2 and -0.2 indicates the threshold for discrimination index (DI) values showing the preference for novel or familiar objects. Absolute DI was plotted to show the magnitude of discrimination regardless of the direction of preference. (A-B) n=12 Con, n=10, IleR n=8, LeuR and n=8 ValR fed 3xTg biologically independent mice. (C) n=8-9 3xTg biologically independent mice per group. (D-E) n=15 Con, n=8 IleR, n=7 LeuR and n=8 ValR fed 3xTg biologically independent mice. (C) n=8 3xTg biologically independent mice per group. (A-F) Statistics for the overall effects of diet, time and the interaction represent the p value from a 2-way ANOVA. *p<0.05, **p<0.01, ***p<0.001, ***p<0.0001, Dunnett’s post-test examining the effect of parameters identified as significant in the two-way ANOVA. Data represented as mean ± SEM.

NOR tests the preference for investigating a familiar object vs a new object and are quantified based on a discrimination index (DI) following a STM and LTM test. The DI is calculated as the difference in time spent exploring the novel versus familiar object, divided by the total exploration time of both objects. A positive DI implies a preference for exploring novelty, indicating that the familiar object’s memory persists and the mice favor exploring the new object. Conversely a negative DI shows that the mice spent more time exploring the familiar object. In both the STM and LTM tests the 3xTg females on IleR and LeuR diets consistently had negative DI; thus these mice failed to explore novel object, and spent more time with the familiar object; ValR-fed females similarly had a negative DI on the STM test (**Fig. 7C**). On the LTM test, ValR-fed females showed a neutral DI, indicating they were unable to discriminate between objects; however, these animals still had significantly greater NOR than Control-fed females (**Fig. 7C**). We also plotted absolute DI to see the overall strength of discrimination irrespective of that being novel or familiar and we did not see any significant changes between any groups in both STM and LTM (**Fig. 7C**).

In contrast to females, LeuR-fed male 3xTg mice showed significant improvements in Barnes maze latency to target during both the STM and LTM tests, and IleR-fed male 3xTg mice showed significantly improved latency to goal during the STM test only (**Fig. 7D**). This was associated with a decrease in errors, with IleR, LeuR, and ValR-fed 3xTg males having fewer errors during both the STM and LTM tests compared to Control-fed mice (**Fig. 7E**). In the NOR test, all the three BCAA restricted groups of 3xTg males had a positive DI that was significantly greater than the negative DI of the Control-fed mice during the STM trial, demonstrating improved recognition memory (**Fig. 7F, left**). In the LTM trial, only IleR-fed males showed a positive DI, which was significantly greater than the Control-fed males, again demonstrating improved memory. When we assessed the magnitude of discrimination by plotting absolute DI, all groups were statistically equivalent (**Fig. 7F, right**).

Finally, after completing the metabolic, behavioral and molecular phenotyping, we next visualized the global effects of individual BCAA restriction on overall phenotypic traits. We performed Principal Component Analysis (PCA) using a combined phenotypic data set including 26 traits including metabolic, behavioral, and pathological data. In 3xTg males, PCA showed distinct clustering between the Control, IleR, LeuR, and ValR groups, with the greatest separation shown between Control-fed and ValR-fed groups (**Figs. S4A-B**). Looking at the key variables, we found hippocampal plaque load, energy expenditure (light and dark), and RER (dark) as some of the strongest contributors to the second principal component (Component 2) driving the separation. In female 3xTg mice, PCA also showed clear separation between the diet groups, particularly between Control-fed mice and mice fed either the ValR or IleR diets (**Figs. S4C**). The strongest contributors to first principal component (Component 1) driving this separation in females were end body weight, lean and fat mass (**Figs. S4C-D**).

### IleR improves the survival of male 3xTg mice

Previous studies, including our own have reported high mortality rates in male 3xTg-AD mice, and we have shown that PR promotes survival of these animals ^15,47,48^. Although female 3xTg mice had low mortality and no differences were observed between diet groups, we similarly observed high mortality of male 3xTg mice in these experiments and observed a significant improvement in survival with IleR (log-rank test, Control vs. IleR, p=0.0354) (**Figs. 8A-B**). ValR- fed mice also trended towards increased survival, while surprisingly LeuR-fed mice had the worst survival outcomes.

**Figure 8:**
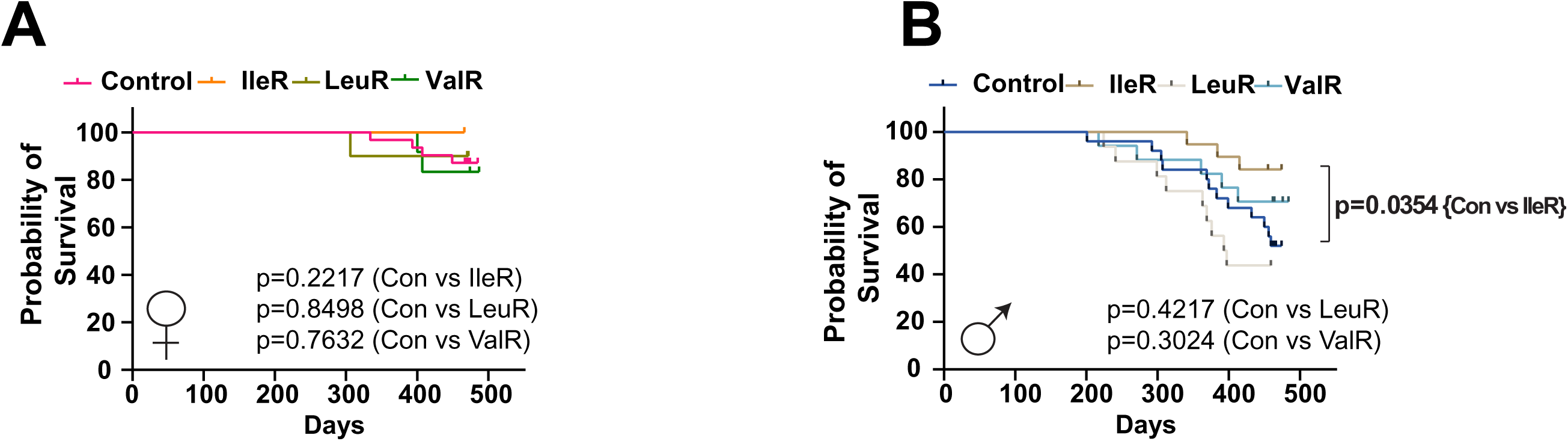
Isoleucine restriction improves survival in male 3xTg-AD mice. (A-B) Kaplan-Meier plots of the survival of female (A) and male (B) 3xTg female mice fed the indicated diets starting at 6 months of age. (A) For females: n=30 Con, n=10 IleR,n=10 LeuR and n=12 ValR fed 3xTg biologically independent mice. (B) For males: n=21 Con, n=20 IleR, n=15 LeuR and n=15 ValR fed 3xTg biologically independent mice. p=0.0345, log-rank test Con vs. IleR mice.

### Gene co-expression network analysis identifies modules associated with metabolic and cognitive traits in males

We next performed Weighted Gene Co-Expression Network Analysis (WGCNA) to generate gene co-expression networks to identify gene modules associated with metabolic, cognitive, and AD-related traits in the brains of female and male 3xTg mice. WGCNA showed several distinct co-expression network modules, each one associated with specific phenotypic traits, including metabolic health, cognitive function, and AD pathology (**Figs. 9A and S5**). These modules were assigned with different colors for classification.

**Figure 9:**
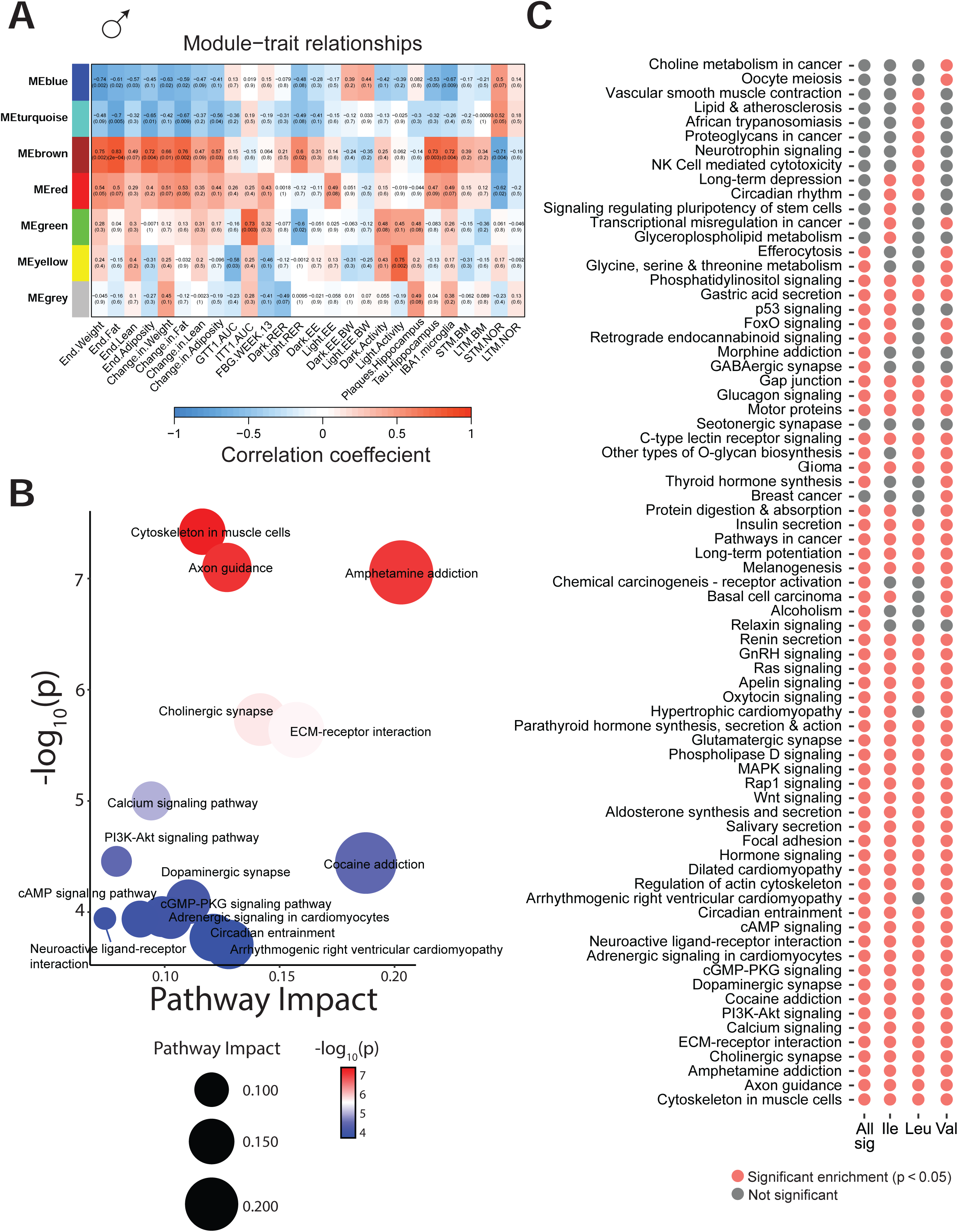
Weighted gene co-expression network analysis identifies modules associated with metabolic, cognitive and pathological outcomes in males. (A) Weighted gene co-expression network analysis (WGCNA) identifies the relationship of module eigengenes (rows) and measures phenotypes (columns) in 3xTg male mice. Heatmap shows the Pearson correlation coefficient between module eigengenes and phenotypic traits, numbers in brackets indicate the corresponding p values. (B) Bubble plot of pathway enrichment based on differentially expressed genes within the brown module following individual BCAA restriction. Red indicates upregulated and blue indicates downregulated pathways relative to Control-fed mice. (C) Dot plot summarizing pathway enrichment for genes in the brown module across all three BCAA restricted diets. Red dot denotes the presence of significant enrichment for a given pathway across all diet groups pooled (All sig) or in the indicated diet group, grey dot denotes absence of significant enrichment.

In males, the highest associations were observed in the Brown module which was significantly correlated with both metabolic and cognitive traits. Genes in this module were positively correlated with increased body weight, fat mass, tau pathology in the hippocampus and glial cell activation and negatively correlated with STM in the NOR test, indicating that metabolic dysfunction and higher tau pathology is linked to worsened cognitive outcomes (**Fig. 9A and Table S5**). The red module also had positive associations with both metabolic parameters and AD pathology and was negatively correlated with STM in the NOR test (**Fig. 9A**). In contrast, the blue and turquoise modules exhibited strong negative correlations with body weight, fat mass, glucose intolerance, tau pathology and glial activation in the hippocampus, and were positively associated with STM performance in the NOR test. In females, WGCNA didn’t identify modules with strong associations with aspects of AD pathology or cognitive performance (**Fig. S5 and Table S6**).

Since the brown module showed the strongest associations with both metabolic and cognitive outcomes, we performed pathway enrichment analysis on the genes within the brown module (**Fig. 9B and Table S7**). Genes in this module were enriched for pathways including axon guidance, cytoskeletal remodeling, and amphetamine addiction, all of which are essential for neuronal connectivity and synaptic plasticity ^49,50^ (**Fig. 9B**). Other pathways enriched in this module included key neurotransmission and intracellular signaling pathways such as dopaminergic synapse, cAMP signaling, and PI3K-Akt signaling (**Fig. 9B**).

We further examined the biological mechanisms associated with the brown module by identifying pathways as defined by the DEGs, first considering all of the BCAA-restricted diets together and then considering IleR, LeuR, and ValR-fed mice separately. The vast majority of these pathways were significantly altered in all of the diets, including cytoskeleton in muscle cells, axon guidance, calcium signaling, oxytocin signaling and MAPK signaling pathway (**Fig. 9C and Table S8**). We also observed pathways that were identified as significantly altered in only one diet group; for example, “Glycerol phosopholipid metabolism” and “p53 signaling” were unique to IleR-fed mice, while “Neurotrophin signaling” and “Natural Killer (NK) cell mediated cytotoxicity” were unique to LeuR-fed mice, and “Choline metabolism in cancer” and “Oocyte meiosis” were altered in ValR-fed mice. To sum up, while there were many similarities in the pathways induced by restriction of any of the three BCAAs, restriction of each BCAA also had distinct transcriptional activity patterns that could mediate many of the BCAA-specific effects we observed on metabolism, AD pathology, and cognition in males.

## Discussion

Dietary protein is a key regulator of metabolic health in rodents and humans. A major contributor to the benefits of a low protein diet is the restriction of branched-chain amino acids (BCAAs), particularly isoleucine, which has been shown to recapitulate many of the metabolic advantages of overall protein restriction ^16^. Restriction of protein, branched-chain amino acids, or isoleucine not only extends lifespan, but promotes favorable molecular changes, rejuvenating the liver transcriptome, improving hepatic aging rate indicators, and reducing hepatic cellular senescence ^12,17,51,52^. Interventions that slow aging should be effective in the prevention or treatment of age-related disease. In support of this hypothesis, we recently demonstrated that PR enhances cognitive function and delays the progression of AD pathology in 3xTg mice, and others have identified benefits of BCAA restriction for AD in this same mouse model ^15,19^.

However, a critical question remained unanswered—do all BCAAs contribute equally to these benefits, or do they have distinct roles in mitigating AD progression? Here, we examined the effects of individually restricting leucine, isoleucine, or valine on AD progression using the 3xTg mouse model of this disease, initiating the study at 6 months of age – an age at which 3xTg mice show cognitive deficits as well as aspects of AD pathology. We discovered that each of the BCAAs has a unique and sex-specific effect on metabolic health, AD neuropathology, and cognitive performance. Strikingly, while ValR significantly improved cognitive function in females, it did not in males; conversely, IleR and LeuR enhanced cognition in males but not in females.

Interestingly, these cognitive improvements occurred independently of reductions in Aβ plaque burden in either of the sexes. Indeed, restricting valine increased the plaque deposition in the hippocampus of female mice, yet ValR was still able to improve memory in female mice relative to females on a Control-fed diet. In fact, ValR had the most pronounced memory improvements in females, further demonstrating that pathological burden may not always align with cognitive functioning. This observation is consistent with findings from clinical trials, where drug candidates targeting amyloid pathways have yielded more modest cognitive improvements than initially hoped despite reducing amyloid burden ^3,53–57^.

Instead, improved cognitive function tended to correlate with reductions in tau pathology. Cognitive improvements following IleR or LeuR occurred in parallel with reduced hippocampal tau phosphorylation, indicating a close alignment with tau pathology and behavioral outcomes in males. While this was not observed in ValR-fed females, which also showed improved cognitive function, these mice did show a reduction in whole brain tau phosphorylation. In our WGCNA analysis of 3xTg males, we found that tau phosphorylation was associated with both worse metabolic outcomes – higher weight and adiposity, worse glycemic control – as well as decreased cognitive function. These findings suggest that mechanisms beyond classical amyloid accumulation that are linked with tau pathology as well as organismal metabolism, may play a key role in mediating the cognitive benefits of BCAA restriction.

Metabolic dysfunction is a hallmark of AD, with insulin resistance and impaired glucose metabolism contributing to neuronal energy deficits ^58,59^. Recent studies have identified elevated plasma BCAA levels, particularly isoleucine, in AD patients, suggesting a possible link between BCAA dysregulation and disease progression ^22,23,60^.Consistent with these findings, our study demonstrates that restricting isoleucine and valine had the most beneficial effects on body weight and adiposity in females and males, whereas leucine restriction had a detrimental impact, leading to increased adiposity in both sexes.

Importantly, our findings offer new mechanistic insights into how BCAA restriction influences AD progression. Despite worsening insulin sensitivity and increased plaque formation, valine restriction emerged as one of the most significant BCAAs in improving cognition in females. Moreover, this improvement in cognition occurred independently of changes in insulin sensitivity, neuroinflammation, mTORC1 signaling, or autophagy activation in the brain. This suggests a potential uncoupling of physiological and molecular pathways from behavioral outcomes, challenging the conventional understanding that cognitive improvements must be directly linked to reductions in AD pathology.

Our findings align with the Cognitive Reserve (CR) hypothesis ^61,62^, which suggests that there can be a mismatch between pathological burden and cognitive performance. This is precisely what we observe in our study, where cognitive improvements occur despite persistent neuropathology. Furthermore, while mTORC1 inhibition and autophagy activation are traditionally linked to neuroprotection, our findings suggest that cognitive improvements can arise through mechanisms distinct from these pathways.

Another interesting observation in our study was that both isoleucine and leucine restricted animals exhibited reduced phosphorylated tau in the hippocampus of both sexes. This suggests that the improvements in cognition observed in males may be partly mediated via tau reductions, and thereby providing evidence that tau pathology may have a stronger impact on cognitive decline than amyloid burden.

Impaired autophagy is a well-recognized contributor to the dysregulation of Aβ metabolism in AD ^63,64^. In our study, IleR led to a reduction in plaque deposition in both males and females, accompanied by decreased microglial activation as shown by reduced IBA1 expression. However, quite surprisingly, most autophagy-related proteins and autophagosome formation markers were downregulated in females from the IleR group with little to no change in males as well. This suggests that a compensatory mechanism, potentially involving the ubiquitin–proteasome system (UPS) or enhanced phagocytic clearance, may have played a significant role under BCAA restriction ^65^. Another possibility is that autophagy was less activated due to a reduced burden of misfolded or aggregated proteins, leading to lower demand for degradation pathways.

In line with these findings, the brain transcriptomic analysis of the brain across all three BCAA-restricted groups identified several distinct and overlapping molecular pathways that correlated with metabolic, behavioral, and neuropathological outcomes. These pathways were mostly observed in males rather than females. Interestingly, we observed many downregulated pathways with very few pathways being upregulated across all groups in 3xTg males. The “Metabolic Pathways” and “Valine, leucine and isoleucine degradation pathways” were upregulated in all the three individual BCAA restricted groups. The upregulation of these pathways might suggest a compensatory metabolic adaptation to reduced BCAA intake for maintaining energy demands.

Limitations of the present study include the exclusive use of the 3xTg mouse model; the use of other AD mouse models could give improved insight, particularly into understanding if the effects of individual BCAAs are mediated by its effects on Aβ, tau, or both. Further, as different strains of mice have different metabolic responses to BCAAs, the effects of BCAA on AD development and progression may vary because of genetic background. Our study focused on a single phosphorylation site of tau, Thr231, previously implicated in AD pathology ^66,67^. However, multiple tau phosphorylation sites contribute to AD pathology, and the effect of BCAA on these sites could be examined in future studies.

Our molecular analyses primarily concentrate on changes in the whole brain; a more extensive and comprehensive analysis will be necessary to elucidate the intricate molecular mechanisms engaged by BCAAs within the different regions of the brain. For instance, analyzing hippocampus and cortex, where we saw dramatic improvements in memory would be beneficial. Additionally, even though we examined autophagy-related proteins, we did not directly assess autophagic flux, which will be beneficial for distinguishing between autophagy initiation and impaired degradation.

In summary, we have shown that restricting each of the BCAA’s have unique and distinct effects in slowing down AD progression in 3xTg mice. Restricting dietary isoleucine and valine broadly improves metabolic health in both the sexes while restricting leucine had negative impacts on overall metabolic health. While restricting isoleucine and leucine were both beneficial in suppressing AD pathology, valine restriction worsened the pathological outcomes primarily in females. Despite this, ValR females exhibited significant cognitive improvements, suggesting that cognitive benefits may occur independently of AD pathology. In contrast, IleR and ValR had a more pronounced effect on memory recognition in males. On a molecular level we observed downregulation of mTORC1 signaling and autophagy in IleR mice. Finally restricting isoleucine improved survival outcomes of male mice. These findings suggest that each BCAA has distinct and sex-specific effects, with significant uncoupling between metabolic, cognitive, and neuropathological outcomes. Taken together, these results provide new insights into the distinct effects of each BCAA in delaying the progression of AD, highlighting BCAA restriction as a promising nutritional approach in slowing or preventing the progression of this devastating disease.

## Materials and Methods

### Animals

All procedures were performed in accordance with institutional guidelines and were approved by the Institutional Animal Care and Use Committee (IACUC) of the William S. Middleton Memorial Veterans Hospital and the University of Wisconsin-Madison IACUC (Madison, WI, USA). Male and female homozygous 3xTg-AD mice were obtained from The Jackson Laboratory (Bar Harbor, ME, USA) and were bred and maintained at the vivarium with food and water available *ad libitum*. Prior to the start of the experiments, at 6 months of age, mice were randomly assigned to different groups based on their body weight and diet. Mice were acclimatized on a 2018 Teklad Global 18% Protein Rodent Diet for 1 week before randomization. Mice were singly housed. All mice were maintained at a temperature of approximately 22°C, and health checks were completed on all mice daily.

At the start of the experiment, mice were randomized to four different groups 1) Control (Amino Acid defined diet), 2) Isoleucine Restricted (IleR), 3) Leucine Restricted (LeuR), and 4) Valine Restricted (ValR) groups; Diet descriptions, compositions and item numbers are provided in **Table S1.**

### In Vivo Procedures

Glucose tolerance test was performed by fasting the mice overnight and then injecting glucose (1 g kg^−1^) intraperitoneally (i.p.) as previously described ^68,69^. For insulin tolerance we fasted the mice for 4 hours and injected insulin intraperitoneally (0.75 U kg^-1^). Glucose measurements were taken using a Bayer Contour blood glucose meter (Bayer, Leverkusen, Germany) and test strips. Mouse body composition was determined using an EchoMRI Body Composition Analyzer (EchoMRI, Houston, TX, USA). For determining metabolic parameters [O2, CO2, food consumption, respiratory exchange ratio (RER), energy expenditure] and activity tracking, the mice were acclimated to housing in an Oxymax/CLAMS-HC metabolic chamber system (Columbus Instruments) for ∼24 h and data from a continuous 24 h period was then recorded and analyzed. Mice were euthanized by cervical dislocation after a 3 hr fast and tissues for molecular analysis were flash-frozen in liquid nitrogen or fixed and prepared as described in the methods below.

### Behavioral assays

All mice underwent behavioral phenotyping at 12 months of age. The Novel object recognition test (NOR) was performed in an open field where the movements of the mouse were recorded via a camera that is mounted above the field. Before each test mice were acclimatized in the behavioral room for 30 minutes and were given a 5 min habituation trial with no objects on the field. This was followed by test phases that consisted of two trials that are 24 hrs apart: Short term memory test (STM and Long-term memory test (LTM). In the first trial, the mice were allowed to explore two identical objects placed diagonally on opposite corners of the field for 5 minutes. Following an hour after the acquisition phase, STM was performed and 24 hrs later, LTM was done by replacing one of the identical objects with a novel object. The results were quantified using a discrimination index (DI), representing the duration of exploration for the novel object compared to the old object.

For Barnes maze, the test involves 3 phases: habituation, acquisition training and the memory test. During habituation, mice were placed in the arena and allowed to freely explore the escape hole, escape box, and the adjacent area for 2 min. Following that during acquisition training the mice were given 180 s to find the escape hole, and if they failed to enter the escape box within that time, they were led to the escape hole. After 4 days of training, on the 5^th^ day (STM) and 12^th^ day (LTM) the mice were given 90 s memory probe trials. The latency to enter the escape hole, distance traveled, and average speed were analyzed using Ethovision XT (Noldus).

### Immunoblotting

Tissue samples from the left hemisphere of the brain were lysed in cold RIPA buffer supplemented with phosphatase inhibitor and protease inhibitor cocktail tablets (Thermo Fisher Scientific, Waltham, MA, USA) using a FastPrep 24 (M.P. Biomedicals, Santa Ana, CA, USA) with bead-beating tubes (16466–042) from (VWR, Radnor, PA, USA) and zirconium ceramic oxide bulk beads (15340159) from (Thermo Fisher Scientific, Waltham, MA, USA). Protein lysates were then centrifuged at 13,300 rpm for 10 min and the supernatant was collected. Protein concentration was determined by Bradford (Pierce Biotechnology, Waltham, MA, USA). 20 μg protein was separated by SDS–PAGE (sodium dodecyl sulfate–polyacrylamide gel electrophoresis) on 8%, 10%, or 16% resolving gels (ThermoFisher Scientific, Waltham, MA, USA) and transferred to PVDF membrane (EMD Millipore, Burlington, MA, USA). The phosphorylation status of mTORC1 substrates p-S240/S244 S6 and 4E-BP1 T37/S46 were assessed in the brain along with autophagy markers including autophagy proteins ATG5, ATG7, and ATG16, as well as autophagosome formation proteins Beclin and light chain 3A/B (LC3A/B), and the autophagy receptor p62 (sequestosome 1, SQSTM1). Tau pathology was assessed by Western blotting with anti-tau antibody. Antibody vendors, catalog numbers and the dilution used are provided in **Table S9.** Imaging was performed using a Bio-Rad Chemidoc MP imaging station (Bio-Rad, Hercules, CA, USA). Quantification was performed by densitometry using NIH ImageJ software.

### Histology for AD neuropathology markers

Mice were euthanized by cervical dislocation after 3 hours fast, and the right hemisphere was fixed in formalin for histology whereas the left hemisphere was snap-frozen for biochemical analysis. For amyloid plaque staining, briefly, brain sections were deparaffinized and rehydrated according to standard protocol. For epitope retrieval, mounted slides were pretreated in 70% formic acid at room temperature for 10 min. Tissue sections were subsequently blocked with normal goat serum (NGS) at room temperature for 1 hr, then incubated with monoclonal antibodies 6E10 (1:100), at 4°C overnight. Aβ immunostained profiles were visualized using diaminobenzidine chromagen. For p-Tau staining and glial activation, brains were analyzed with anti-GFAP (astrocytic marker), and anti-Iba1 (microglial marker) antibodies respectively. The following primary antibodies were used: phospho-Tau (Thr231) monoclonal antibody (AT180) (Thermo Fisher Scientific; # MN1040, 1:100) anti-GFAP (Thermo Fisher; # PIMA512023; 1:1,000), anti-IBA1 (Abcam; #ab178847; 1:1,000). Sections were imaged using an EVOS microscope (Thermo Fisher Scientific Inc., Waltham, MA, USA) at a magnification of 4X, 10X and 40X magnification. Image-J was used for quantification by converting images into binary images via an intensity threshold and positive area was quantified.

### Transcriptomic Analysis

RNA was extracted from the left hemisphere of the brain using trizol followed by Purelink RNA mini kit (cat#12183018A; Thermo Fisher Scientific, Waltham, MA). The concentration and purity of RNA was determined using a NanoDrop 2000c spectrophotometer (Thermo Fisher Scientific, Waltham, MA) and RNA was diluted to 100–400 ng/µl for sequencing. The RNA was then submitted to the University of Wisconsin-Madison Biotechnology Center Gene Expression Center & DNA Sequencing Facility, and RNA quality was assayed using an Agilent RNA NanoChip. RNA libraries were prepared using the TruSeq Stranded Total RNA Sample Preparation protocol (Illumina, San Diego, CA) with 250ng of mRNA, and cleanup was done using RNA Clean beads (lot #17225200). Reads were aligned to the mouse (Mus musculus) with genome-build GRCm38.p5 of accession NCBI:GCA_000001635.7 and expected counts were generated with ensembl gene IDs74.

Analysis of significantly differentially expressed genes (DEGs) was completed in R version 3.4.3 ^70^ using *edgeR* ^71^ and *limma* ^72^. Gene names were converted to gene symbol and Entrez ID formats using the mygene package. Male and female mice were analyzed separately, and clear outliers were removed following PCA analysis of the raw data. Genes with too many missing values were removed, if genes were present in less than one diet/age group they were removed. To reduce the impact of external factors not of biological interest that may affect expression, data was normalized to ensure the expression distributions of each sample are within a similar range. We normalized using the trimmed mean of M-values (TMM), which scales to library size. Heteroscedasticity was accounted for using the voom function, DEGs were identified using an empirical Bayes moderated linear model, and log coefficients and Benjamini-Hochberg (BH) adjusted p-values were generated for each comparison of interest ^73^. Pathway enrichment was conducted using KEGG pathways on significant genes for each BCAA restricted group vs Control (BH Adjusted p<0.05) using the *limma* package.

WGCNA analysis was conducted in R (Version 4.4.3) using the WGCNA package ^74^. First, we filtered by gene expression variance, keeping the top 50% variable genes to remove “noisy” genes, this reduced gene number from 15524 to 7762 genes (males) and from 17468 to 8734 (females). We then checked that all genes had enough samples and there were no clear outliers before running the analysis. WGCNA analysis identifies significant modules of genes and their associations with phenotypes. Once modules of genes were identified, they were enriched for KEGG Pathways.

### Statistical Analysis

All statistical analyses were conducted using Prism, version 9.0.2 (GraphPad Software Inc., San Diego, CA, USA). Tests involving multiple factors were analyzed by either a two-way analysis of variance (ANOVA) with diet and time or sex as variables or by one-way ANOVA, followed by a Dunnett’s post-hoc test as specified in the figure legends. Alpha was set at 5% (p < .05 considered to be significant). Kaplan–Meir survival analysis of 3xTg mice was performed with log-rank comparisons stratified by sex and diet. Data are presented as the mean ± SEM unless otherwise specified.

## ACKNOWLEDGEMENTS

We thank all the members of the Lamming lab for their feedback. The Lamming lab is supported in part by the NIA (AG056771, AG062328, AG081482, and AG084156), the NIDDK (DK125859), by a grant from the Alzheimer’s Association (23AARG-1029665), and by startup funds from UW-Madison. RB was supported in part by F31AG081115. CLG was supported in part by Dalio Philanthropies, a Glenn Foundation for Medical Research Postdoctoral Fellowship, and by grant HF-AGE AGE-009 from the Hevolution Foundation to CLG. MFC was supported in part by F31AG082504. CYY is supported in part by K99AG084921. The Puglielli lab is supported in part by the NINDS (NS094154), the NIGMS (GM148487) and the NIA (AG078794). The authors thank the University of Wisconsin Carbone Cancer Center Experimental Animal Pathology Laboratory supported by P30 CA014520, for use of its facilities and services. LP and DWL are members of the Wisconsin Nathan Shock Center of Excellence in the Basic Biology of Aging, P30 AG09258601. The Lamming lab was supported in part by the U.S. Department of Veterans Affairs (I01-BX004031 and IS1-BX005524), and this work was supported using facilities and resources from the William S. Middleton Memorial Veterans Hospital. The content is solely the responsibility of the authors and does not necessarily represent the official views of the NIH. This work does not represent the views of the Department of Veterans Affairs or the United States Government.

## DATA AVAILABILITY

RNA-sequencing data have been deposited with the Gene Expression Omnibus and are available under accession number GSE299928. The authors declare that source data supporting the findings of this study are available within the paper and its supplementary information and Source Data files.

## AUTHOR CONTRIBUTIONS

RB, MMS, FX, CLG, MFC, MT, AT, RM, CYY, IG, SS, BK conducted the experiments. RB, CLG, DV and DWL analyzed the data. RB, MJB, LP and DWL wrote and edited the manuscript.

