## Supplementary Figures and Table Legends for "Restriction of individual branched-chain amino acids has distinct effects on the development and progression of Alzheimer’s disease in 3xTg mice"

### Supplementary Figure 1

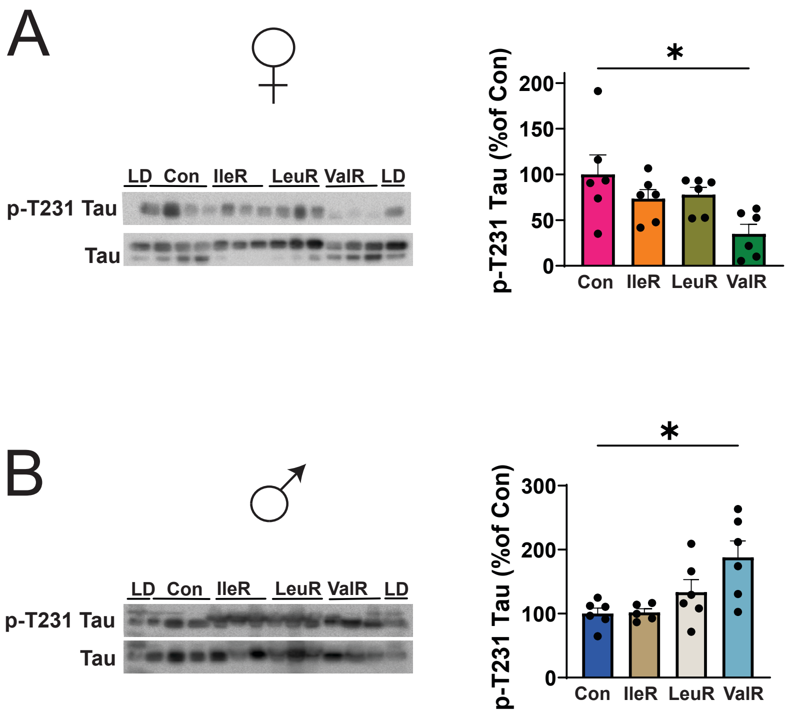

#### Supplementary Figure Legends

##### Supplementary Figure 1: Individual BCAA restriction effects on Tau phosphorylation in the whole brain of 3xTg mice

Western blot analysis of phosphorylated T231 Tau in whole brain lysates of (A) female and male (B) 3xTg mice. (A-B) n=6 3xTg biologically independent mice per group. *$p<0.05$, **$p<0.01$, ***$p<0.001$, ****$p<0.0001$ Dunnett's post-test examining the effect of parameters identified as significant in the one-way ANOVA. Data represented as mean ± SEM.

### Supplementary Figure 2

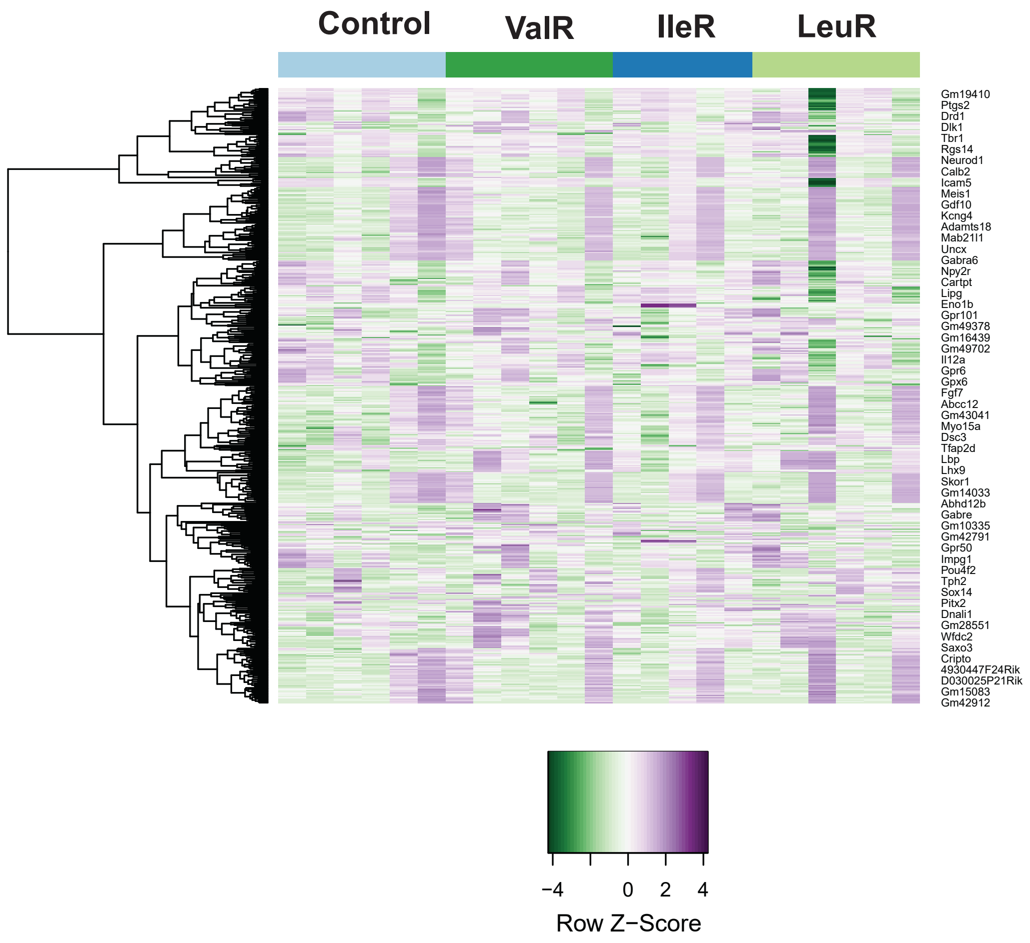

**Supplementary Figure 2: Individual BCAA restriction did not impact transcripts in female brain.**

Heatmap of the top 50 differentially expressed (DEG) genes in females. n=5-6 animals/group.

### Supplementary Figure 3

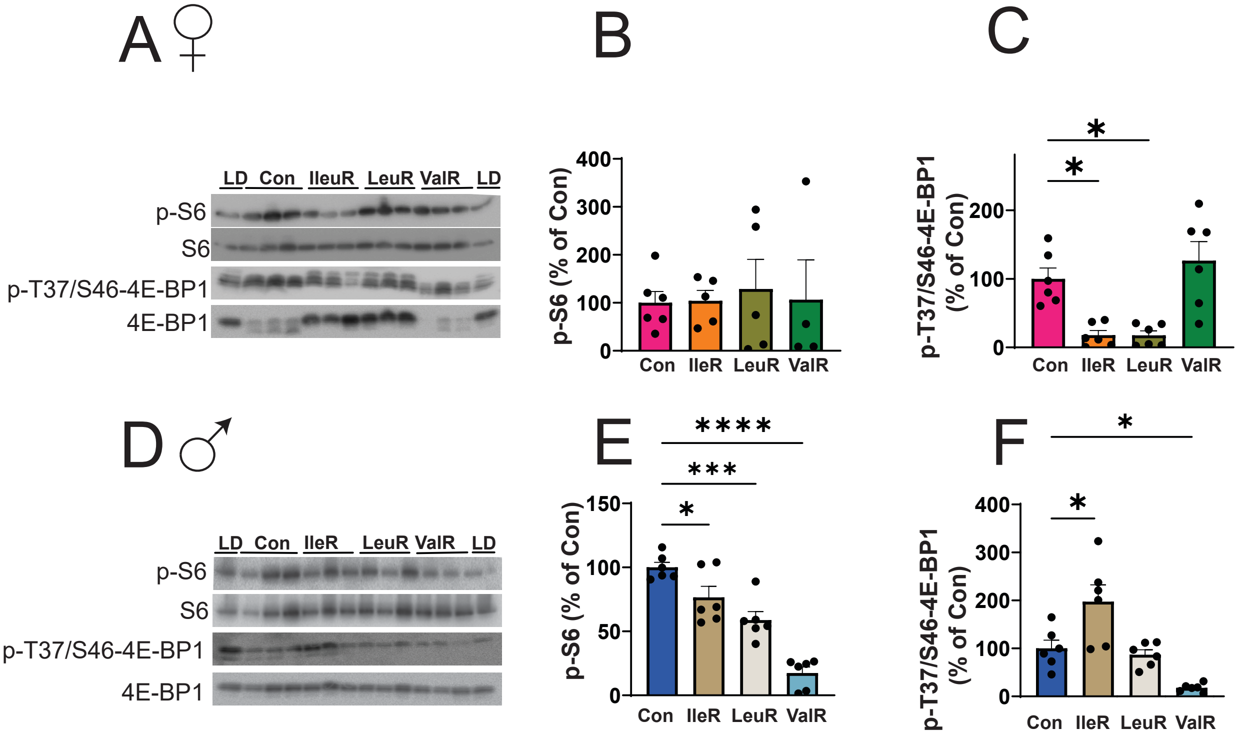

**Supplementary Figure 3: mTORC1 signaling in the brain of 3xTg mice following individual BCAA restriction**

(A-F) Western blotting of mTORC1 signaling substrates in the whole brain lysates of 3xTg mice. (A, D) Representative immunoblot of p-S240/S244 S6 and T37/S46 4E-BP1 in females (A) and males (D). (B, E) Quantification of the phosphorylation of p-S240/S244 S6 (B) in females and (E) males relative to expression of S6 in (C, F) Quantification of the phosphorylation of T37/S46 4E-BP1 in (C) females and (F) males relative to expression of 4E-BP1. *$p<0.05$, ***$p<0.001$, ****$p<0.0001$ Dunnett's post-test examining the effect of parameters identified as significant in the one-way ANOVA. Data represented as mean ± SEM.

### Supplementary Figure 4

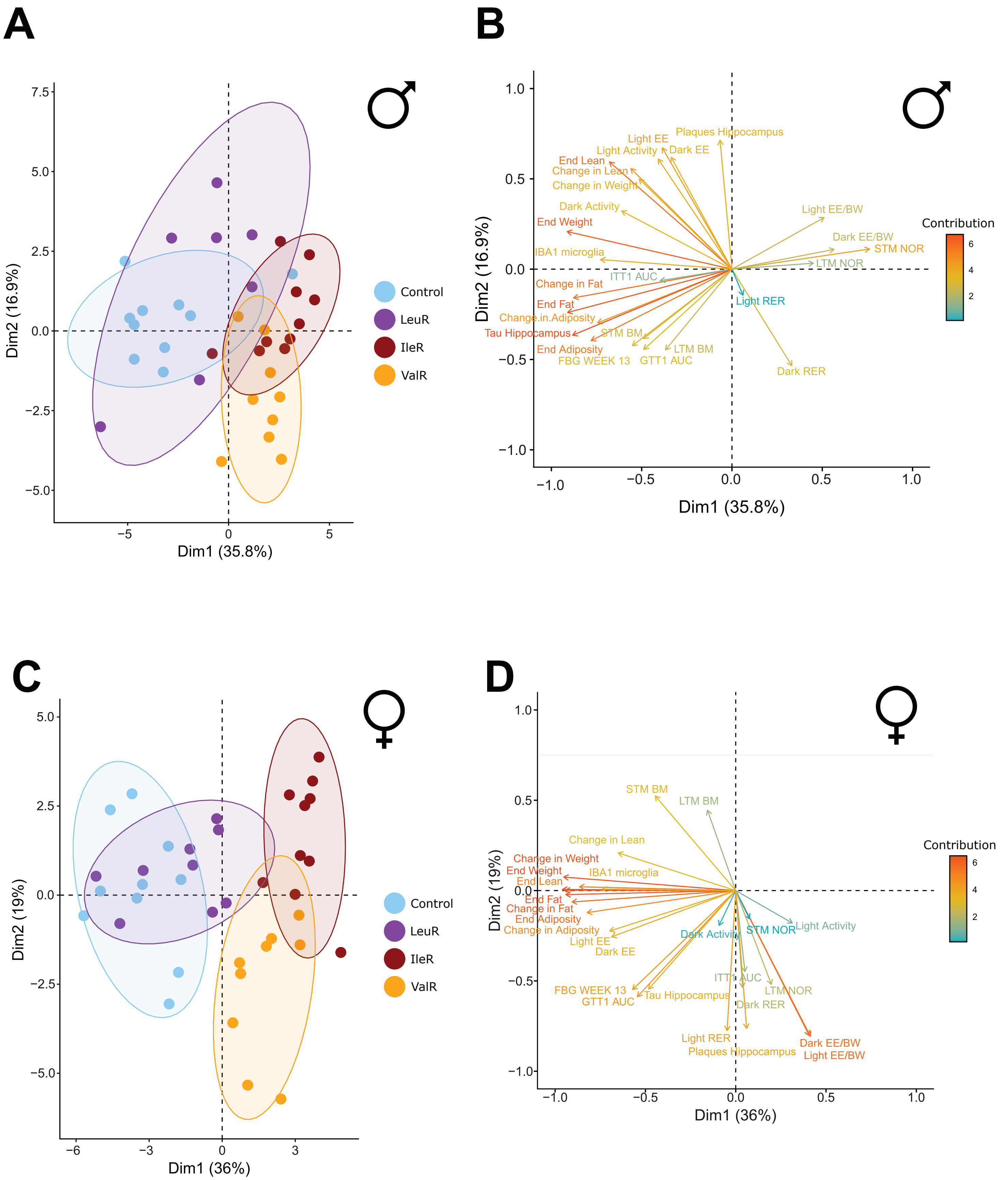

**Supplementary Figure 4: Principal component analysis of combined phenotypic traits in both sexes.**

Principal component analysis of the phenotypic traits and the variables contributing to the PCA spread in (A-B) in males and (C-D) in females.

### Supplementary Figure 5

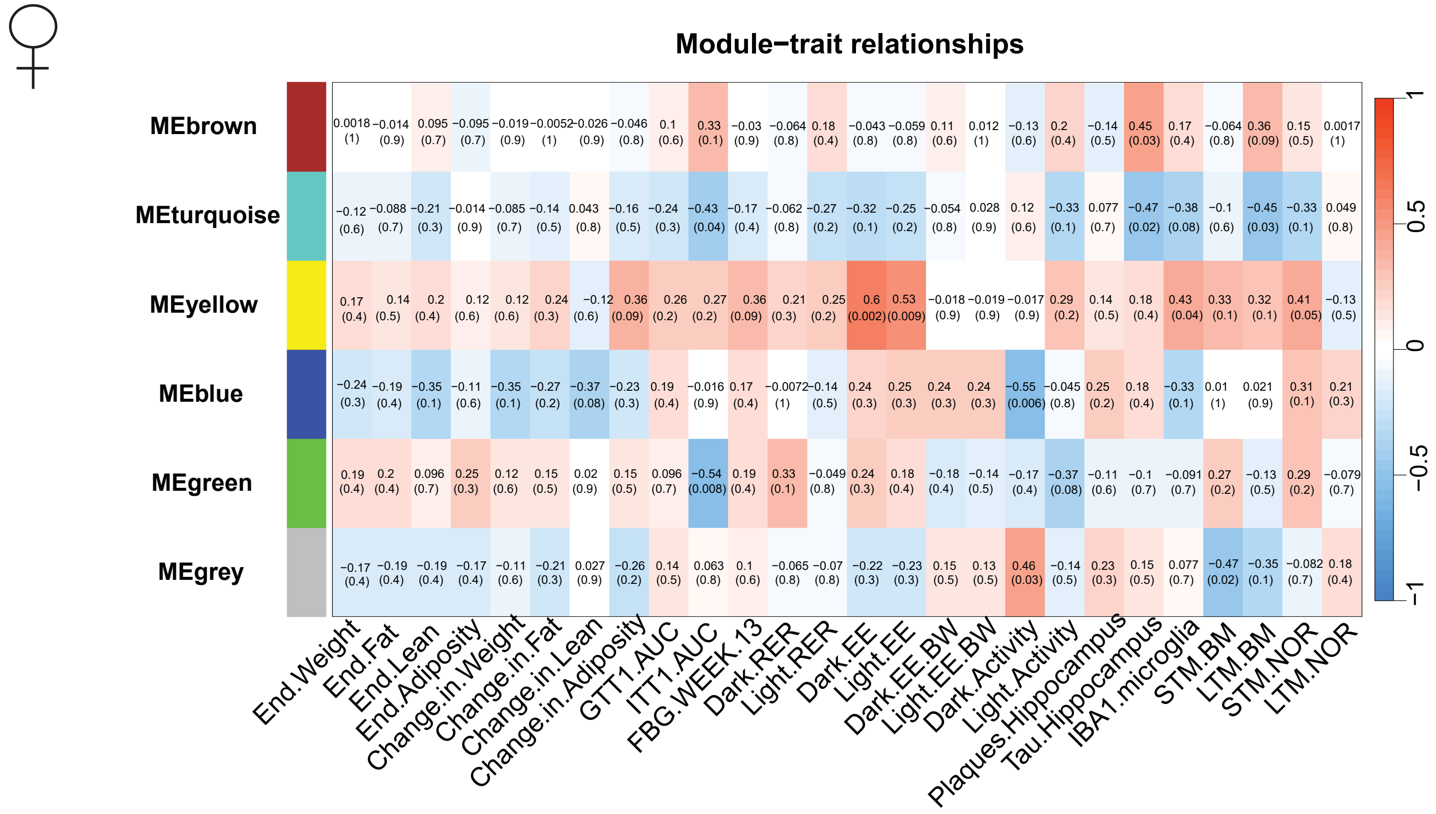

**Supplementary Figure 5: Gene co-expression network analysis identifies modules associated with metabolic, cognitive and pathological outcomes in females.**

Weighted gene co-expression network analysis (WGCNA) identifies the relationship of module eigengenes (rows) and measures phenotypes (columns) in 3xTg female mice. Heatmap shows the Pearson correlation coefficient between module eigengenes and phenotypic traits, numbers in brackets indicate the corresponding p values.

**Supplementary Table Legends**

**Supplementary Table 1:** Diet composition and calorie content for diets used in this study.

**Supplementary Table 2:** Differentially expressed gene names, $log_2$ fold-changes and related p-values from transcriptomic analysis in the brains of male 3xTg mice across diet groups.

**Supplementary Table 3:** Differentially expressed gene names, $log_2$ fold-changes and related p-values from transcriptomic analysis in the brains of female 3xTg mice across diet groups.

**Supplementary Table 4:** KEGG enriched pathways for significant genes identified in 3xTg male mice as shown in **Figure 5C.**

**Supplementary Table 5:** Gene co-expression modules identified using Weighted Gene Co-Expression Network Analysis (WGCNA) in male 3xTg mice as shown in **Figure 9A.**

**Supplementary Table 6:** Gene co-expression modules identified using Weighted Gene Co-Expression Network Analysis (WGCNA) in female 3xTg mice as shown in **Supplementary figure 5.**

**Supplementary Table 7:** Pathway enrichment analysis of the genes within the brown module in male 3xTg mice as shown in **Figure 9B.**

**Supplementary Table 8:** KEGG pathway enrichment analysis of differentially expressed genes in female 3xTg mice.

**Supplementary Table 9:** Antibodies used for both western blotting and immunohistochemistry.
